# Reduced PABPN1 levels causes cytoskeleton disorganization and aberrant differentiation

**DOI:** 10.1101/2020.01.15.907311

**Authors:** Cyriel Sebastiaan Olie, Erik van der Wal, Cikes Domagoj, Loes Maton, Jessica C. de Greef, I-Hsuan Lin, Yi-Fan Chen, Elsayad Kareem, Josef M. Penninger, Benedikt M. Kessler, Vered Raz

## Abstract

The polyadenylation binding protein nucleus 1 (PABPN1), a multifactorial regulator of mRNA processing, regulates muscle wasting and atrophy. Previously, we elucidated the PABPN1-dependent proteome and found that levels of structural proteins, sarcomeric and cytoskeletal, were highly altered. We identified MURC, a plasma membrane-associated protein, to be affected by the cytoskeletal stability and suggest that MURC is a novel marker for impaired regeneration in muscles. We also studied the spatial organization of muscle structural proteins in 2D and 3D cell models with reduced PABPN1 levels (named here as shPAB). We show that dysregulation of cytoskeletal proteins in the shPab proteome is associated with a cytoskeleton lacking a polarized organization in muscle cells. We show that consequently, the cell mechanical features as well as myogenic differentiation are significantly reduced. We then show that restoring cytoskeletal stability, by actin overexpression in shPAB was beneficial for cell fusion and for the expression of sarcomeric proteins in shPAB models. We suggest that poor cytoskeleton mechanical features are caused by altered expression levels and contribute to aging-associated muscle wasting and atrophy.

## Introduction

Muscle aging and many muscular dystrophies are characterized by reduced muscle strength, atrophy and impaired regeneration capacity (Schiaffino, Dyar et al., 2013). Muscle aging is associated with multiple cellular mechanisms affecting protein homeostasis, transcription, protein acetylation and various metabolic and energy pathways (Anisimova, Alexandrov et al., 2018, Ohlendieck, 2011, Olie, Riaz et al., 2019, Ubaida-Mohien, Lyashkov et al., 2019). As such multiple proteins regulate aging. Reduced levels of polyadenylation binding protein 1 (PABPN1) lead to aging-associated muscle pathology, including atrophy, regeneration, extracellular matrix thickening, and switches in myofiber typing (Riaz, Raz et al., 2016). The cellular mechanisms for how reduced PABPN1 levels leads to muscle atrophy is not fully understood.

PABPN1 is a multifactorial regulator of mRNA processing (Banerjee, Apponi et al., 2013), and in cells with reduced PABPN1 levels RNA expression profiles are altered genome-wide (Abbassi-Daloii, Yousefi et al., 2017, de Klerk, Venema et al., 2012, Jenal, Elkon et al., 2012). In human, PABPN1 levels are reduced from midlife onwards, specifically in skeletal muscles (Anvar, Raz et al., 2013). An expansion mutation in PABPN1 is the genetic cause for oculopharyngeal muscular dystrophy (OPMD), a late onset myopathy (reviewed in: Brais, 2009). In OPMD, reduced levels of functional PABPN1 are caused by both aggregation of the expanded PABPN1 and reduced PABPN1 mRNA levels (Raz & Raz, 2014). Reduced PABPN1 levels (named shPab in mouse and shPAB in human) cause muscle wasting and atrophy (Olie et al., 2019, Riaz et al., 2016).

To elucidate the cellular mechanisms that are affected by reduced PABPN1 levels, proteomic studies were generated in mouse muscles, indicating the possible affected cellular processes (Banerjee, Phillips et al., 2019, Olie et al., 2019). Yet, the cellular mechanisms for how reduced PABPN1 levels leads to muscle atrophy is not fully elucidated. Consistent with reduced muscle strength in aging and in OPMD, sarcomeric and cytoskeletal protein groups were significantly altered in the shPab proteome (Olie et al., 2019). The cytoskeleton provides structure and strength to the cell, and together both protein groups facilitate muscle contraction (Gautel & Djinović-Carugo, 2016). Disruptions in cytoskeletal arrangement, often due to mutations in cytoskeletal proteins, can cause various muscular dystrophies (Spence, Chen et al., 2002). The actin filaments are an essential part in the contractile force of muscles (Pollard & Cooper, 2009). In muscle cell culture, both microtubule and actin filaments have been suggested to be essential for cell fusion (Abmayr & Pavlath, 2012, Mian, Pierre-Louis et al., 2012). It has been proposed that a cross-talk between the actin-cytoskeleton and microtubules facilitates proper organization of the cytoskeleton (Dogterom & Koenderink, 2019). Although disruption of actin organization in aged muscles are prominent (Higuchi-Sanabria, Paul Rd et al., 2018), it is not fully understood how it affects myogenesis. Here we investigated the PABPN1-dependent cytoskeletal changes using 2D and 3D models, and discuss how these affect muscle wasting.

## Methods

### Mouse muscle tissue handling

All mouse muscles that were used in this study have previously been described by C. Olie et al. (Olie et al., 2019). In brief, muscles denoted as shPab or Control are tibialis anterior (TA) muscles injected with AAV9 particles expressing shRNA to *Pabpn1* or scrambled shRNA, respectively, as detailed by M. Riaz et al. (Riaz et al., 2016). AAV9 shPab and scrambled particles were injected contralateral into the left or the right leg, allowing paired analysis. This experimental design overcomes natural variations between mice. Five weeks after AAV9 injection muscles were harvested and directly frozen in liquid nitrogen. From each muscle cryosections were collected for protein analysis (mass-spectrometry and western blot) as previously described (Olie et al., 2019).

### Human muscle tissue

Human vastus lateralis muscles were collected as waste material during a knee operation from healthy individuals with an informed consent (Anvar et al., 2013). Cryosections of 16 µm thick were made with a cryostat CM3050S (Leica Microsystems) and pasted on Super Frost Plus glass slides (Menzel-Glaser; ThermoFisher Scientific).

### Muscle cell culture

Immortalized human muscle cell (7304.1) shPAB and control stable cell lines were generated as previously described (Anvar et al., 2013). In brief, PABPN1 knockdown (shPAB) was generated after lentivirus transduction with a shRNA to *PABPN1*. The control cell line was transduced with scrambled shRNA (Anvar et al., 2013). Myoblast cells were maintained in growth medium (F10 medium supplemented with 15% FCS, 1 ng/ml bFGF, 10 ng/ml EGF and 0.4 ug/ml Dexamethasone). Differentiation (cell fusion) was conducted in DMEM supplemented with 2% horse-serum for 4 days. Stable actin-eGFP overexpressing cells were generated with a lentivirus transduction in control or shPAB cell cultures. The actin-eGFP was subcloned into a lentivirus expression cassette. Actin-eGFP expression was driven by the CMV promoter. All cultures were mycoplasma-free. Microtubule destabilization or stabilization treatments were performed using Nocodazole (250 nM, Cayman chemicals) or Paclitaxel (25 nM, Cayman chemicals), respectively for 2 hours.

### Muscle bundle cultures

Muscle bundles were generated as previously described by Bakooshli et al with minor adjustments (Afshar Bakooshli, Lippmann et al., 2019). Before generation of muscle bundles, bone-shaped PDMS chambers were sterilized and pre-treated with 1% Pluronic® F-127 (Sigma) for 1 hour at room temperature. Confluent myoblast cultures were dissociated with TrypLE™ Express Enzyme (Gibco) reagent and 3×10^5 cells were used per muscle bundle. Cells were incubated on ice and combined with an ice-cold hydrogel mixture consisting of 4 mg/ml bovine fibrinogen (Sigma) and 20% Growth Factor Reduced Matrigel® (Corning). Before loading the cell/hydrogel mixture into a bone-shaped PDMS chamber, 0.8 units of Bovine Thrombin (Sigma) were added to initiate fibrinogen polymerization. Loaded PDMS chambers were incubated for 30 minutes at 37°C and after polymerization muscle bundles were kept in normal proliferation medium containing 1.5 mg/ml 6-aminocaproic acid (6-ACA) (Sigma). After 2 days, differentiation was induced by changing medium to differentiation medium (DMEM high glucose (Gibco), 2% horse serum (Gibco), 1% Penicillin G (Sigma) and 2 mg/ml 6-Aca). Seven days after differentiation muscle bundles were fixed overnight at 4°C with 2% PFA. Muscle bundles were stored in PBS at 4°C.

### Protein analysis

Proteins from TA muscles were extracted as previously described (Olie et al., 2019). Proteins from human muscle cell culture were extracted using RIPA extraction buffer (20 mM Tris, pH 7.4, 150 mM NaCl, 5 mM EDTA, 1% NP40, 5% glycerol and 1 mM DTT and protease inhibitor cocktail). The extracts were briefly sonicated and protein concentrations were determined using Bio-Rad protein assay. Cellular fractionation was done as described by S. Baghirova et al. (Baghirova, Hughes et al., 2015) with some minor adjustments. In brief, cells were trypsinized and were subsequently treated with Nocodazole or DMSO (as control) for 30 minutes. Lysis buffer A (300mM NaCl, 100mM HEPES pH 7.4, 50 ug/mL Digitonin, 2M Hexylene glycol and protease inhibitor cocktail) was directly added after the treatment and incubated for 10 minutes at 4°C. Centrifugation separated the soluble proteins (cytosolic proteins) from the insoluble proteins that remain in the pellet. Insoluble proteins were then collected by the addition of 1% Igepal to lysis buffer A.

Protein extracts were separated using precasted SDS-PAGE (Criterion XT, Bio-Rad). Western blot was carried out using PVDF membranes and blotted with the following antibodies: Actin (sc-8432; Santa Cruz), CSRP3 (GTX110536, GeneTex), GAPDH (MA5-15738 / Pierce), MURC (HPA021021, Atlas antibodies), PABPN1 (LS-B8482, LS Bio, USA) and Tubulin (T6199; Sigma-Aldrich). Primary antibodies were labeled using fluorescent IRDye 800CW or IRDye 680RD (LiCOR) and were subsequently detected using the Odyssey CLx Infrared imaging system (LiCOR, NE. USA). Protein expression levels were quantified using ImageJ version 1.48 (https://imagej.nih.gov/ij/). All expression levels were corrected for background and normalized to loading controls.

### Immunofluorescence

Immunostaining on muscle sections was carried out with the previously described protocol (Olie et al., 2019) using the following primary antibodies; CSRP3 (GTX110536, GeneTex), Dystrophin (ab7164, Abcam), MURC (HPA021021, Atlas antibodies) and specific myosin heavy chain isotype were stained with MyHC-2a and MyHC-2b conjugated to 594 and 488 fluorophore, respectively (Riaz, Raz et al., 2015). Secondary antibodies used were anti-rabbit conjugated to Cy5 or anti-mouse conjugated with Alexa 488 (ThermoFisher). Immunofluorescence in cell cultures was carried out after fixation in 2% formaldehyde and permeabilization in 1% triton-X100. Primary antibodies used for immunofluorescent staining in cell culture were Actin (sc-8432; Santa Cruz), MURC (HPA021021, Atlas antibodies) and Tubulin (T6199; Sigma-Aldrich). Fused cells (myotubes) were visualized using an antibody against Myosin heavy chain isoforms (MyHC) (MF20, Sigma-Aldrich). Primary antibodies were visualized with anti-rabbit or anti-mouse conjugated with Alexa 488 or Cy5 (ThermoFisher). DAPI was used to stain nuclei in both muscle tissue and cell culture. Immunofluorescence of muscle bundles was carried out with the following protocol: After washing with PBS muscle bundles were incubated in a blocking buffer (3% BSA, 0.3% Triton X-100, 0.1% Tween-20 in PBS) for 1 hour and washed in PBS. An incubation with the primary antibody was carried out for one hour in antibody buffer (0.1% BSA, 0.3% Triton X-100, 0.1% Tween-20 in PBS). After PBT washing, secondary antibody mix containing DAPI was incubated for 30 minutes in antibody buffer. After PBT washing muscle bundles were imaged with the Dragonfly 2-photon imaging system; ANDOR Oxford instruments.

### Cell mechanical properties

Probing of cell mechanical properties was carried out with the Brillouin Light Scattering Microscopy (BLSM) and the Atomic Force Microscopy (AFM). A day before analysis, cells were seeded on Matrigel Coated 35mm glass bottom microwell dishes (MatTek Corporation) in growth media at a density of 60-50%. During imaging, samples were kept at 37°C and 5% CO_2_ levels via an onstage heater.

The BLSM was performed using a home built Brillouin confocal microscope described in (Elsayad, Werner et al., 2016). Excitation was via a single mode 532nm laser (Torus, Laser Quantum, DE). For all scans, the laser power at the sample was between 1-3 mW, and the swell time per point, which was also the acquisition time of each spectra, was 100ms. The spectral projection was measured on a cooled EM CCD camera (ImageEMX II, Hamamatsu, JP). The spectrometer was coupled to an inverted microscope frame (IX73, Olympus, JP) via a physical pinhole to assure confocal detection. Spectral phasor analysis (Elsayad, 2019) was used to obtain initial parameter estimates for peak positions and widths which were subsequently inserted into a non-linear least squares fitting algorithm that fitted two broadened Lorentzian functions (Voigt functions) to obtain the two peak positions, from which the Brillouin Frequency Shift (BFS) could be calculated. The BFS is directly proportional to the (longitudinal) acoustic phonon speed, which has been empirically found to be higher for stiffer samples and smaller for softer samples. All data analysis was performed in Matlab (Mathworks, DE) using custom written scripts (Elsayad et al., 2016). The AFM was performed on a JPK nanowizard 4 system. Applied imaging mode used was QI (quantitative imaging) where a force curve is applied at each point. Area analyzed per each cell was 5 µm × 5 µm (64×64 pixels) with an approach speed of 35microns/s (3.4ms/pixel), and applied forces of up to 118pN. The cantilevers used for sample probing (qp-BioAC, CB3, Nanosensors) were calibrated to 828 microNewton/m as a force constant. All data analysis was performed in JPK SPM data processing software.

### Imaging and image quantification

Imaging of the mouse and human sections was carried with the DM5500 fluorescent microscope (Leica Microsystems). Images were captured with the Leica application suite X version: 3.3.3.16958. Mean fluorescence intensity (MFI) was measured from over 400 myofibers and a Pearson correlation was calculated in paired analysis. Immunofluorescence cell staining was imaged using cell-based imaging and subsequent quantification was carried out with the colocalization toolbox in the Arrayscan VTI HCA, Cellomics (Thermo Fisher Scientific). In brief, nuclei were segmented from the DAPI signal and myotubes with the MyHC signal. Cell fusion was calculated by the average fraction of nuclei within MyHC objects.

## Results

### Reduced PABPN1 expression levels highly affect structural proteins

To study structural changes in aging-associated muscle atrophy, we first compared fold-change direction of cytoskeletal genes in the tibialis anterior (TA) muscles of aged mice (Lin, Chang et al., 2018) to the proteome of mouse TA muscles with reduced PABPN1 levels (shPab) (Fig. S1). From the 65 annotated cytoskeletal genes, which were found in both studies, 57% (37/65) showed a similar fold-change direction (Fig. S1). Consistent with the comparison between mRNA and protein fold-changes from our previous study in (Olie et al., 2019), also in this subset of genes we found that the protein fold-changes exceeded the corresponding mRNA fold-changes (Fig. S1). Among the cytoskeletal proteins, tubulin and actin showed similar fold-change directions in both aging and shPab muscles (Fig. S1). Actin levels were slightly down-regulated whereas tubulin levels were up-regulated. This suggests that cytoskeleton organization could be affected in shPAB muscle cells. Paired analysis of protein abundance and western blot validation were carried out for selected proteins. Levels of tubulin, CSRP3 and MURC were up-regulated in shPab muscles (Fig. 1A-B and Fig. S2). Actin levels showed a trend with a small average negative fold-change in shPab (Fig. 1A). CSRP3 and MURC fold changes in both shPab proteome and in aged muscles were significant and >1.5 folds (Fig. S1), and both were reported to be involved in muscle biology (Tagawa, Ueyama et al., 2008, Vafiadaki, Arvanitis et al., 2015). Their involvement in muscle aging and muscle wasting was not investigated before.

**Fig 1.**
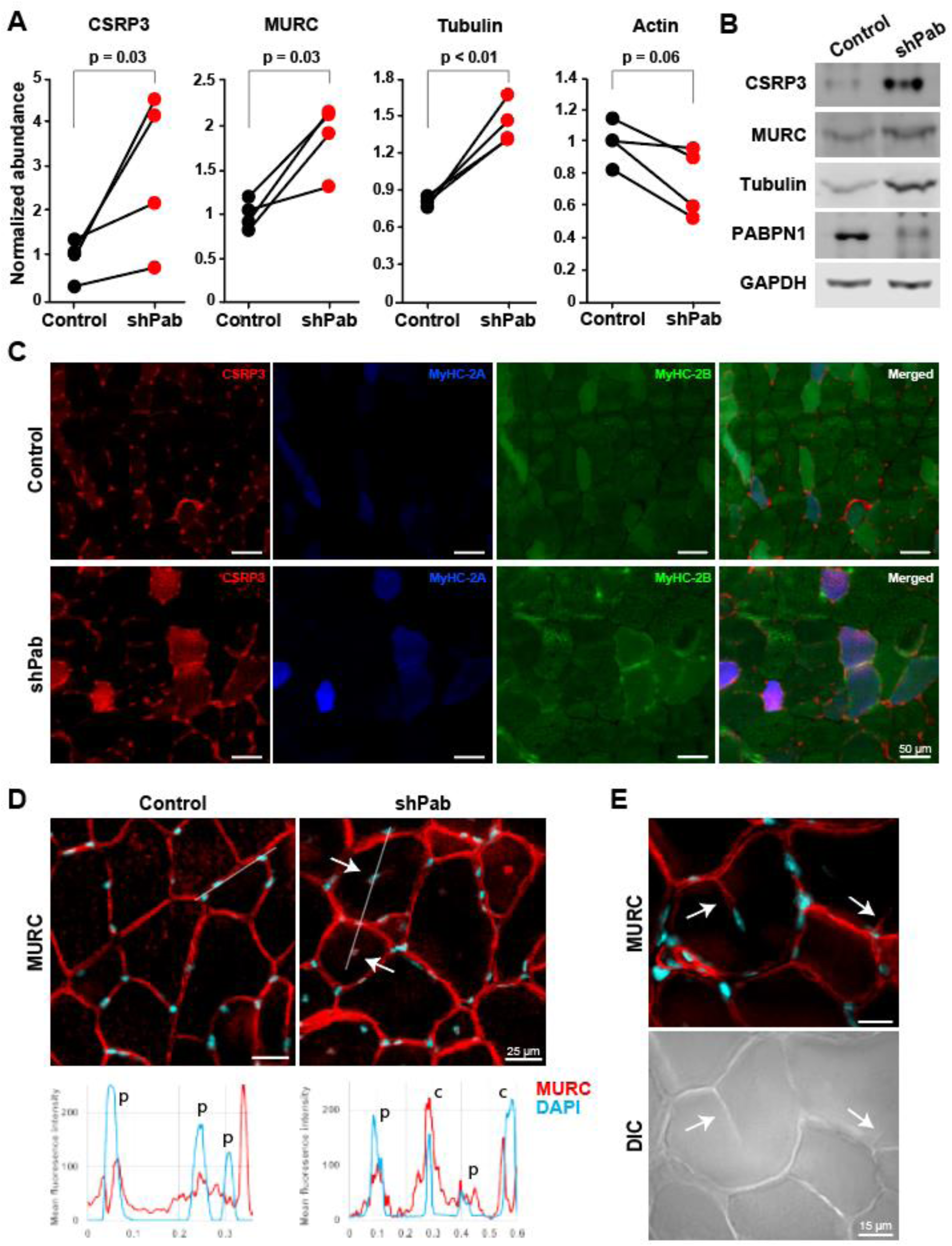
Expression levels and spatial expression of CSRP3 and MURC in shPab muscles. **A.** Paired dot-plots show normalized mass spectrometry abundances of CSRP3, MURC, tubulin, and actin in control and shPab muscles. A statistical difference was assessed with a paired ratio t-test. (N=4 mice). **B.** Western blot shows expression levels in paired muscles. GAPDH was used as loading control. PABPN1 level confirms down-regulation. **C.** Spatial localization of CSRP3 in scram and shPab muscles. Images are from a representative region in TA muscle, showing transitions of MyHC-2B and MyHC-2A transitions between scram and shPab. Staining shows CSRP3 in red, MyHC isoforms type-2A in blue and type-2B in green. The merged image shows CSRP3 enrichment in type-2A myofibers. Scale bar is 50 µm. **D.** Spatial localization of MURC in shPab muscles. Upper panel shows representative images in TA muscles: MURC is depicted in red and DAPI nuclear counterstaining in cyan. MURC staining in central nuclei is shown in the fluorescence distribution plot. Arrows point to central nuclei. Scale bar is 25 µm. Lower panel shows intensity distribution of MURC and DAPI along the white line (in the upper panel) peripheral nuclei and depicted with P, and central nuclei with C. **E.** MURC signal in a myofiber splitting initiation (left, depicted by arrows) in shPab muscle. The same regions are indicated by arrows in a DIC image (right). Scale bar is 15 µm.

Histopathological changes in shPab muscles include myofiber transitions, reduced myofiber size (atrophy), central nuclei and splitting myofibers (Olie et al., 2019, Riaz et al., 2016). We then assessed if those changes involved higher expression levels of CSRP3 or MURC. Muscle cross-sections were stained with an antibody mix to myosin heavy chain (MyHC) isoforms: -2A and -2B and to CSRP3 or MURC. We found that CSRP3 colocalized with MyHC-2A positive myofibers (Fig. 1C and Fig. S3). A correlation of the mean fluorescence intensity (MFI) in individual myofibers showed that CSRP3 MFI correlated with MyHC-2A MFI but not with MyHC-2B MFI (Table S1). The correlation between MyHC-2A and CSRP3 was higher in shPab compared to control muscles (Table S1, r=0.68; r=0.53, respectively). This observation is consistent with higher CSRP3 levels and higher proportions of MyHC-2A positive myofibers in shPab muscles (Riaz et al., 2016), suggesting that higher CSRP3 in shPab muscle is associated with myofiber transition. In contrast, MURC colocalization with MyHC-isoforms was not obvious (Fig. S4). We therefore did not assess a correlation between MURC and MyHC-isoforms in myofibers.

Since shPab muscles were also marked with central nuclei (Olie et al., 2019, Riaz et al., 2016), we then assessed if CSRP3 or MURC up-regulation co-localizes with central nuclei. CSRP3 expression was not found in central nuclei (Fig. S5). MURC staining, however, showed a spatial overlap with DAPI in central nuclei (Fig. 1D). Interestingly, MURC staining was also found to co-localize with a newly formed membrane in between myofibers (Fig. 1E). This suggests that MURC is expressed in regions of splitting myofibers, which were predominantly found in shPab muscles (Fig. 1E).

Regions of splitting myofibers are enriched in injured muscles (Murach, Dungan et al., 2019). To verify MURC spatial expression in regions of splitting myofibers, we stained human muscles from injured young donors. In injured regions, but not in healthy regions, we found MURC to be localized with central nuclei and in splitting myofibers (Fig. 2). The plasma membrane was stained with dystrophin, and MURC co-localized with dystrophin (Fig. 2). This can suggest that MURC expression is associated with muscle regeneration. Consistently, in shPab muscles PAX7 expression in myonuclei was increased compared with control (Olie et al., 2019). An increase in Pax7 and in splitting myofibers suggests an increase in regeneration due to reduced PABPN1 levels.

**Fig 2.**
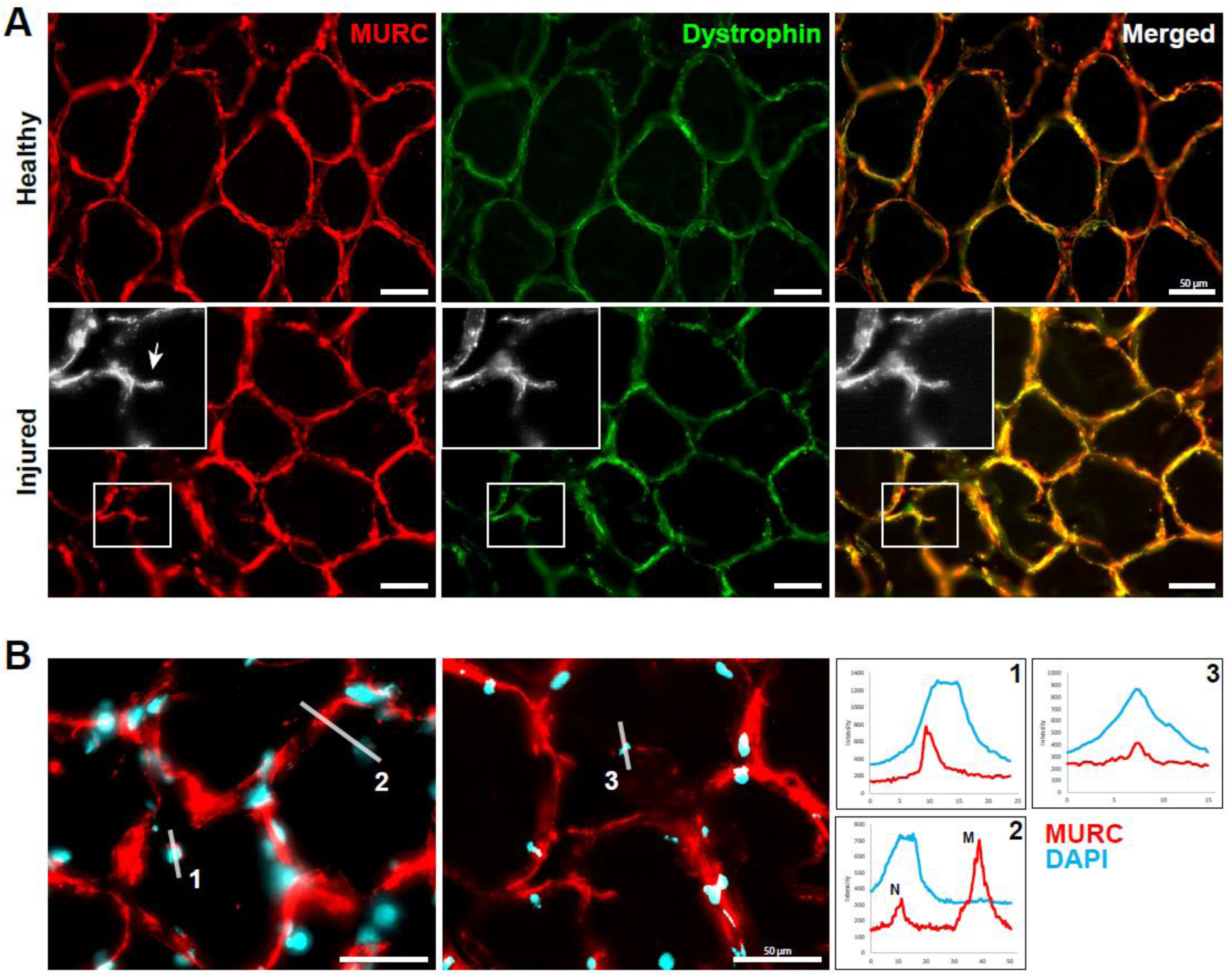
Cytoskeletal changes in human skeletal muscles are also characterized by MURC expression. **A.** Immunohistochemistry of human muscle cross-sections. Images represent healthy or injured regions. MURC is stained in red and dystrophin is stained in green, an overlap is in the merged image. A myofiber splitting, pointed by an arrow, is shown in the enlarged grayscale image, and the corresponding region is highlighted with a square. Scale bar is 50 µm. **B.** MURC staining in central nuclei in wounded muscles is shown in two muscles. MURC is depicted in red, and nuclei with cyan. For three regions (marked 1-3 with a white line), a corresponding intensity distribution plot is shown in the right panel. MURC is in red, and DAPI in cyan. Scale bar is 50 µm.

### MURC localization marks the aberrant cell fusion in shPAB cultures

To elucidate a role for MURC in PABPN1-mediated myogenic defects, we studied MURC expression in a stable shPAB human muscle cell culture. MURC expression levels in human myoblast culture were low compared to myotube cell culture (Fig. 3A), which is in agreement with previous studies (Bastiani, Liu et al., 2009, Tagawa et al., 2008). MURC immunofluorescence staining in myoblasts showed a predominant localization in foci, one per cell (Fig. 3B). MURC foci were found to colocalize with tubulin foci (Fig. 3B). In mitotic cells, tubulin foci were reported to be associated with centrosomes, playing a role in microtubule organization and spindle formation in mitotic cells (Baumgart, Kirchner et al., 2019, Woodruff, Ferreira Gomes et al., 2017). This suggests that MURC is also associated with the centriole. MURC foci in shPAB myoblasts were significantly larger (Fig. 3C). Moreover, in both control and shPAB myoblasts we found the MURC foci to actually be two foci, but in shPAB myoblasts MURC foci were more distant from each other (Fig. 3D-E).

**Fig 3.**
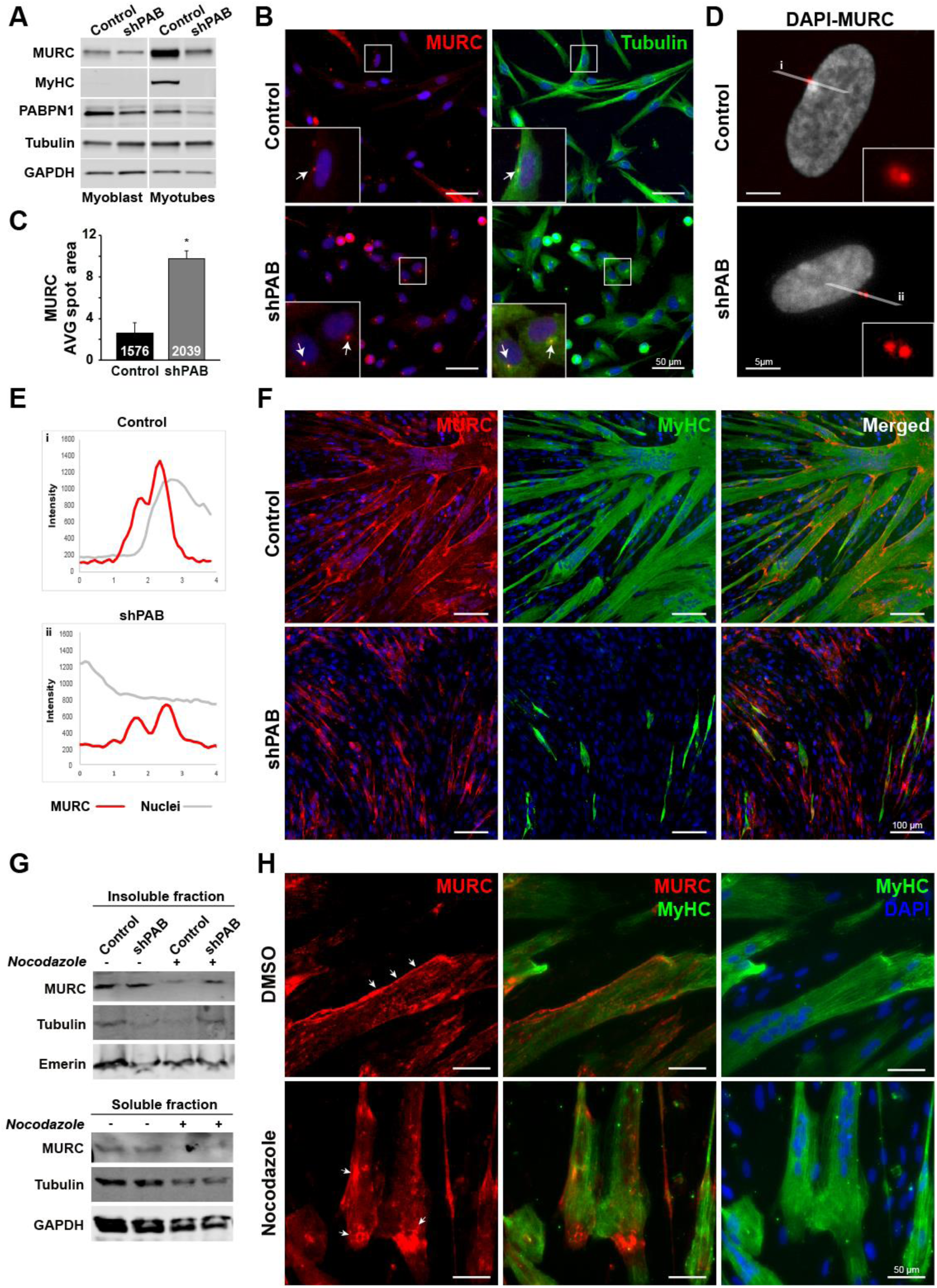
MURC localization in shPAB human muscle cell cultures. **A.** Western blot shows MURC expression levels in myoblast and myotubes, respectively. GAPDH was used as loading control. PABPN1 level confirms down-regulation. **B.** Representative images of control and shPAB human muscle cell cultures. Control and shPAB myoblasts are stained with tubulin (green) and MURC (red). White arrows point to MURC foci and the colocalization with tubulin. Nuclei are counterstained in blue with DAPI. Scale bar is 50 µm. **C.** Bar chart shows average MURC foci area in control and shPAB cultures. Numbers inside bars depict the number of nuclei, averages and standard deviations are from (N=3). Statistical significance was assessed with the Student’s t-test **D.** Images show MURC foci in red and nuclei are counterstained with DAPI in white. Grey lines (i or ii) indicate the region of which the MFI overlay is shown in **E. F.** Control and shPAB fused cultures are stained for MyHC (green) and MURC (red). Merged images show overlay between MyHC and MURC. A nuclear counterstaining is in blue. Scale bar is 100 µm. **G.** A western blot of MURC in soluble and insoluble fractions in control and shPAB cell cultures. Nocodazole treatment was carried out in protein extract. Nocodazole treatment is monitored by tubulin levels. Emerin and GAPDH are loading controls for the insoluble or soluble fractions, respectively. **H.** Images show MURC (in red) localization in mock-treated and nocodazole-treated myotubes (depicted with MyHC in green). White arrows indicate MURC localization at the plasma membrane. Scale bar is 50 µm.

In fused cultures MURC levels were elevated in both control and shPAB compared to myoblast cultures (Fig. 3A). Control myotubes had high MyHC expression that went together with high MURC expression, whereas shPAB fused cultures had no myosin expression but still expressed more MURC compared to control shPAB myoblasts (Fig. 3A). In control cultures MURC expression was enriched in fused myotubes (Fig. 3F). In contrast, in shPAB cell culture, MURC signal was not co-localized to MyHC expressing cells (Fig. 3F). Moreover, in control myotubes MURC outlines the myotube membrane but not in shPAB cells (Fig. 3F). MURC was previously described to be associated with caveolin at the cell plasma membrane (Naito, Ogata et al., 2015). A western blot analysis showed MURC enrichment in the insoluble fraction in both control and shPAB cell cultures (Fig. 3G). This observation confirmed MURC plasma membrane localization in control fused cultures, but not in shPAB culture. The membrane localization in shPAB cells could suggest that MURC trafficking to the plasma membrane is abolished. Trafficking and protein association to the plasma membrane can be stabilized by the microtubule (Stephens, 2012). Therefore, we then investigated if MURC plasma membrane localization is affected by microtubule stability. Treatment with nocodazole, a β-tubulin binding compound that depolarizes microtubules, disrupted MURC localization at the plasma membrane in fused cell cultures (Fig 3H). Moreover, MURC levels in the insoluble fraction were also reduced by nocodazole treatment (Fig. 3G). In contrast, insoluble MURC levels in shPAB cell culture were unchanged by nocodazole treatment (Fig. 3G). This suggests that MURC dis-trafficking to the plasma membrane in shPAB is associated with aberrant microtubule organization.

### Reduced PABPN1 levels in muscles cells lead to an aberrant cytoskeleton architecture and reduced cell tension

Cells with reduced PABPN1 levels are characterized by reduced myogenesis (Anvar et al., 2013, Apponi, Leung et al., 2010), but the cellular causes have not been investigated. As dynamic cytoskeletal reorganization plays a role in myogenesis (Guerin & Kramer, 2009), we investigated the cytoskeleton organization in shPAB cell culture and whether myogenesis in shPAB cells could be enhanced by modulation of the cytoskeleton. In control myoblasts both tubulin and actin immunofluorescence staining showed elongated filaments in the longitudinal cell axis (Fig. 4A), but in the shPAB myoblasts structured filaments were not visible and the cells were more rounded (Fig. 4A). We then visualized actin filaments with an overexpression of actin-GFP. Whereas the actin filaments were stretched and parallel organized along the longitudinal axis in the control cells, in shPAB cells actin filaments were disorganized and unstructured (Fig. 4A). A rounded cell shape was also observed after myoblasts treatment with drugs that modulate microtubule stability (Fig. 4B). Together, this suggests that polarized cytoskeletal organization was not formed or maintained in shPAB myoblasts, which directly influences biophysical cell properties. We then measured the mechanical cell properties in both control and shPAB cell cultures using two different methods. The Brillouin Light Scattering Microscopy (BLSM) measures the cell hydrostatic pressure and cytoplasm viscoelasticity, which was found to be lower in shPAB cells (Fig. 4Ci). The Atomic Force Microscopy (AFM) measures cell surface tension, which was also found to be significantly lower in shPAB cells (Fig. 4Cii). With the AFM larger differences and smaller variations were found because the measurements are more focal compared with the BLSM. In both methods reduced values were found in shPAB, confirming each other. Reduced stiffness and reduced hydrostatic pressure and reduced cytoplasm viscoelasticity are consistent with the observed reduced cytoskeletal filament organization in shPAB cells.

**Fig 4.**
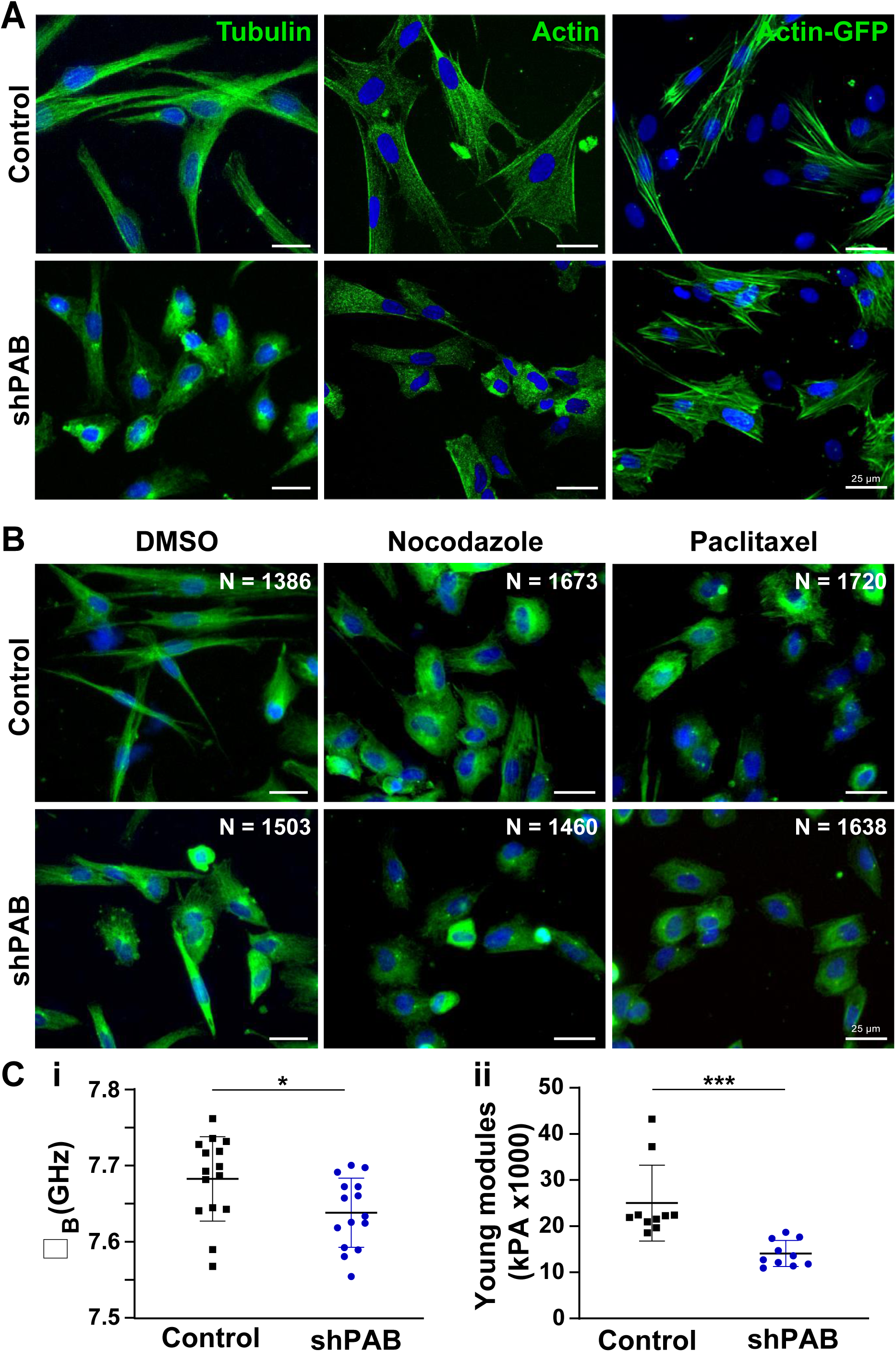
Disrupted cytoskeletal spatial organization in shPAB human muscle cell cultures. **A.** Representative images of control and shPAB human muscle cell cultures, stained with antibodies to tubulin and actin, and the actin filaments were visualized with actin-GFP. **B.** Tubulin staining in control and shPAB myoblast cell cultures after DMSO, 100 nM nocodazole or 25 nM paclitaxel treatment for 2 hours. Scale bar is 25 µm. **C.** Measurements of cell-mechanics properties in control and shPAB cells using the Brillouin Light Scattering Microscopy (BLSM; Ci) or the Atomic Force Microscopy (AFM; Cii). Measurements were carried out in myoblasts; every dot represents the median from 1000 measurements in a cell. Cell stiffness is measured by GHz, and the young modulus reports the cell surface tension. Averages and standard deviations are from N=15 cells. Statistical significance was calculated with the student’s t-test.

We then investigated if reduced cytoskeletal organization affect cell fusion in human muscle cells. To test this, myoblasts were treated with drugs affecting microtubule stability and were subsequently incubated with fusion medium for four days. As described above, nocodazole depolarizes microtubules by binding β-tubulin. In contrast to this destabilizing property of nocodazole, Paclitaxel supports microtubule stability by binding to the αβ-tubulin heterodimer. Paclitaxel treatment during myogenic differentiation led to reduced cell number and consequently to a reduced fusion index in both control and shPAB cell cultures (Fig. 5A). Moreover, after paclitaxel treatment the MyHC-positive cells were narrower and contained fewer nuclei compared to untreated control myotubes (Fig. 5A). A two-hour treatment with nocodazole, a microtubule destabilizing agent, had no obvious effect on fused cells or on the fusion index (Fig. 5A). As the effect of nocodazole is reversible (Musa, Orton et al., 2003, Samson, Donoso et al., 1979), this suggests that microtubule destabilization does not play a role in the first phase of cell fusion. Together, this data suggests that during the first phase of cell fusion, destabilization of microtubule filaments but not stabilization, might play a role.

**Fig 5.**
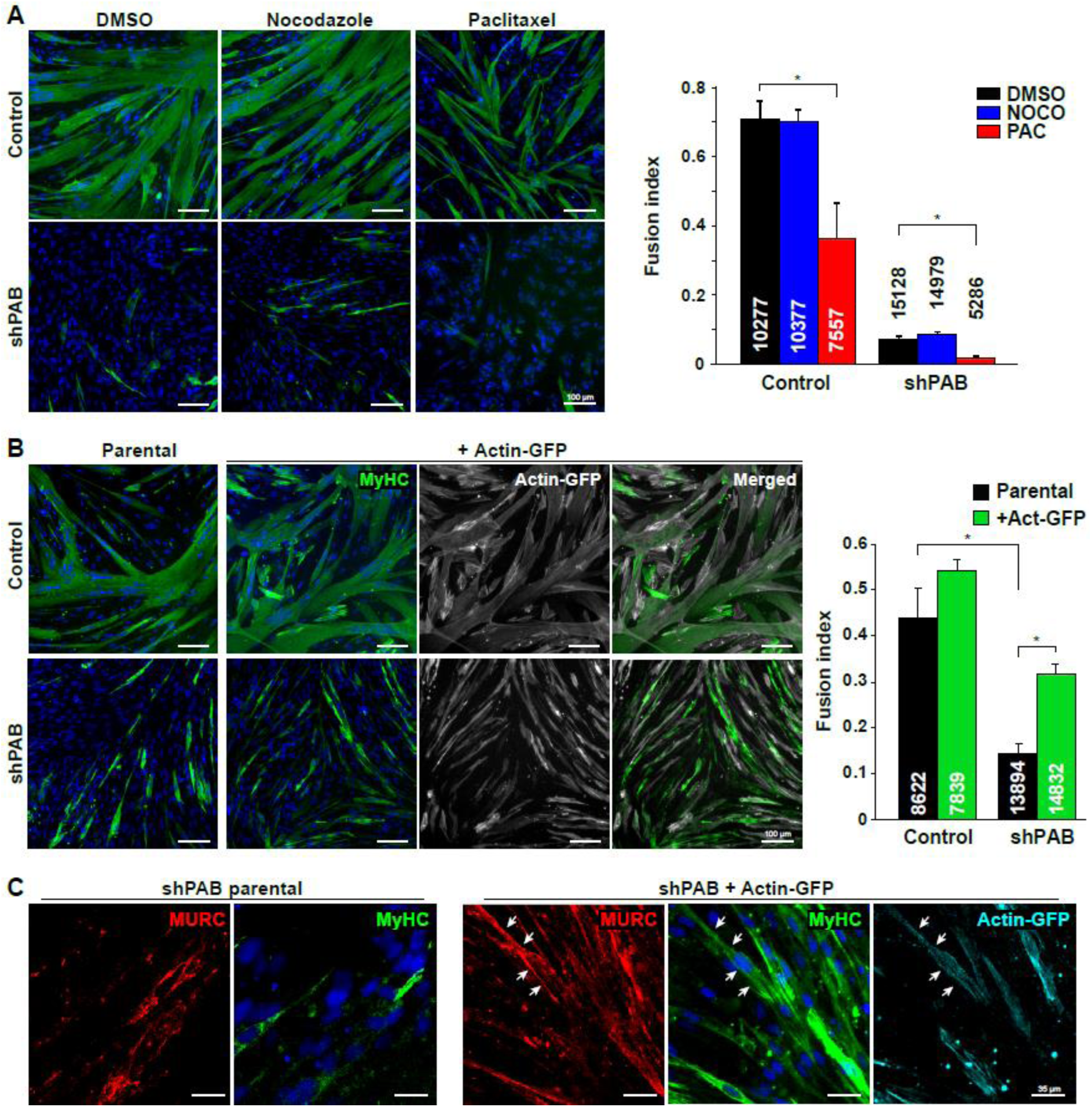
Reduced myogenesis in shPAB is affected by cytoskeleton stabilization and organization. **A.** Representative images of fused control and shPAB cultures. MyHC staining is depicted in green and the nuclei in blue. Bar chart shows fusion index, which was calculated by the fraction of nuclei within MyHC objects. The average number of nuclei per replicate is denoted in or above each bar. Average and standard deviation are from four replicates. Scale bar is 100 µm. **B.** Cell fusion in the actin-GFP cell lines. Left panel: Representative images of cell cultures fused for four days, parental or actin-GFP overexpressed control or shPAB cell lines. MyHC is depicted in green and actin-GFP in gray, the merged image shows an overlay between MyHC and actin-GFP. Bar chart (left panel) shows fusion index in parental and actin-GFP control and shPAB cultures. Average and standard deviation are from three replicates. The average number of nuclei per replicate is denoted in each bar. Scale bar is 100 µm. **C.** Images show MURC localization (depicted in red) in the actin-GFP (depicted in cyan) fused cell cultures. Scale bars is 35 µm. Statistical significance was assessed using a student’s T-test, * denotes p<0.05. Every experimental setup was repeated three times.

To further investigate the hypothesis that cytoskeletal organization affects myogenesis in shPAB cells, we studied the ability to fuse in control or shPAB cell lines that stably express actin fused to GFP. The percentage of GFP-positive cells was similar between control and shPAB cultures (Fig. S6), ensuring that any differences in cell fusion were not due to differences in transduction efficiency. In control cells, actin-GFP expression did not affect fusion index (Fig. 5B). In contrast, MyHC expression in shPAB cell culture significantly increased by actin-GFP expression (Fig. 5B). However, those MyHC-expressing cells were not multinucleated and were narrower compared to control myotubes (Fig. 5B). We then further characterized the myotubes in the shPAB+actin-GFP expressing cell culture by studying MURC expression. MURC signal was found to colocalize with the membranes of MyHC-expressing cells in shPAB+actin-GFP cell culture (Fig. 5C). This colocalization to membranes in fused cultures was absent in the parental shPAB cultures (Fig. 5C). Together, this data suggests that stabilizing the actin filaments could enhance myogenesis and restore MURC localization at the plasma membrane in shPAB cell culture. However, stabilization of actin filaments is not sufficient to form multinucleated myotubes in a 2D cell culture system.

To validate and expand on the 2D cell culture results, we then employed the 3D muscle model, which has a better maturation of fused cells. We first assessed if muscle bundle could be generated from the shPAB muscle cells and then whether actin overexpression would be beneficial. After seven days of differentiation, muscle bundles were formed from both parental and shPAB cell cultures (Fig. 6A). Consistent with the 2D cell culture, the muscle bundles from control culture expressed MyHC (Fig. 6B). In the shPAB muscle bundle the expression of MyHC was clearly reduced compared with control (Fig. 6B and 6C, respectively). As the same fusion medium was used in both 2D and 3D cultures, but to the 3D muscle bundle a synthetic matrix was added, it suggests that cell fusion in the shPAB muscle bundle fusion is triggered by the synthetic matrix. Nevertheless, those muscle bundles may not form a contractible unit, as they express low levels of the MyHC isoforms. We then investigated muscle bundles of shPAB+actin-GFP, and MyHC was clearly expressed (Fig. 6C). We then compared MURC expression in shPAB parental cells to shPAB+actin-GFP, and consistent with the 2D cell model, in shPAB+actin-GFP muscle bundle MURC expression was clearly increased compared with shPAB parental (Fig. 6C). MURC staining in shPAB+actin-GFP was found in elongated cells (Fig. 6C). Together, the data suggests that actin overexpression is beneficial for expression of sarcomeric genes in shPAB cultures and partly restores the MURC expression.

**Fig 6.**
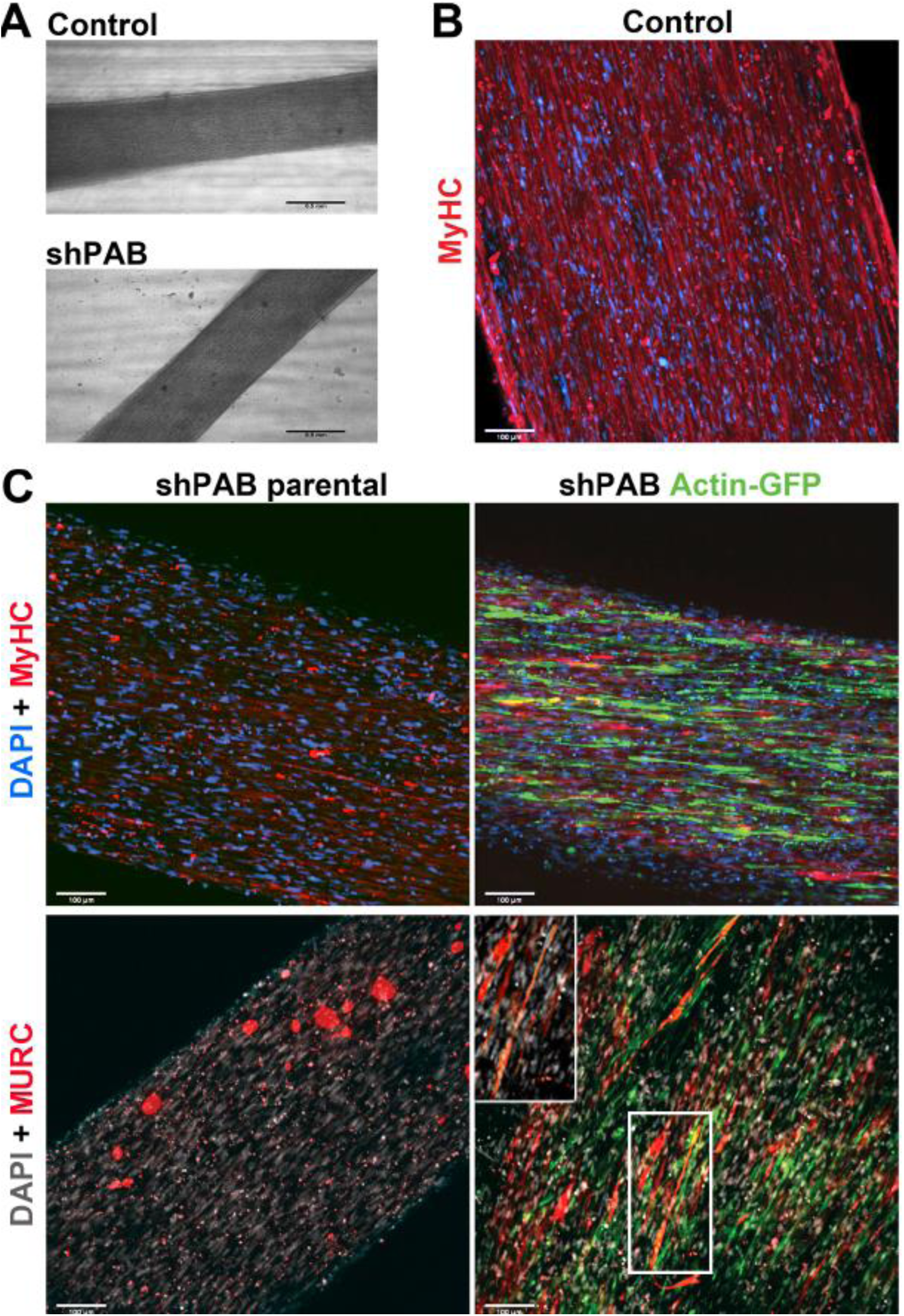
shPAB cultures form muscle bundles that lack myosin heavy chain expression. **A.** Bright field images of muscle bundles generated from control and shPAB muscle cells. Scale bar is 500 µm. **B.** MyHC staining in muscle bundle from control cells. **C.** Maximum projection of muscle bundle staining using shPAB parental and shPAB+actin-GFP cells. MyHC or MURC are in red, GFP is in green. Nuclei are counterstained with Hoechst. Scale bar is 100 µm. Insert shows a magnification of MURC staining in multinucleated cells.

## Discussion

Reduced PABPN1 levels cause muscle atrophy (Riaz et al., 2016). Atrophic muscles show alterations in cytoskeleton organization, which is not fully understood (Schiaffino, 2017). We found significant changes in expression levels of cytoskeletal proteins in shPab muscles. Fold-change direction for some of those proteins was found to be in common with muscle aging. We show that in cell cultures with stable PABPN1 down-regulation the cytoskeleton structures, microtubules and actin filaments lack orientation and organization. The small and round morphology of shPAB myoblasts implies an inability to obtain an elongated and polarized muscle cell shape. Both altered expression levels of cytoskeletal proteins and a disorganized architecture of the cytoskeletal filaments would influence the cell mechanical properties. Consistently, we found that both cell stiffness and cell surface tension are reduced in shPAB myoblasts. The defects in both cellular and mechanical features lead to a failure to start the fusion as the cytoskeleton (Kim, Jin et al., 2015).

Disrupted microtubule structures have been observed in muscular dystrophies such as Duchenne muscular dystrophy (DMD) (Khairallah, Shi et al., 2012). In a recent study, disrupted microtubule organization was found to be associated with higher levels of tubulin proteins (Randazzo, Khalique et al., 2019). We also found elevated levels of tubulin proteins in the shPab proteome, including β-tubulin6 (Tubb6) (Olie et al., 2019). Tubb6 knockdown was found to be beneficial for microtubule organization in the DMD mouse model (Randazzo et al., 2019). Moreover, Tubb6 levels were higher in myofibers with central nuclei, suggesting a role in regeneration (Randazzo et al., 2019). Consistently, markers of regeneration are higher in shPab muscles (Olie et al., 2019), and we show that cytoskeleton disorganization in shPAB 2D cell culture is causally linked to reduced myogenesis. Microtubule disorganization negatively affects cell fusion (Mian et al., 2012). Targeting of microtubule stability, using small molecules drugs, was not beneficial for cell fusion (Mian et al., 2012, Musa et al., 2003). Similarly, treatments affecting microtubule stability were not beneficial for cell fusion in shPAB cell cultures.

Consistent with a recent study showing that muscle aging in C. elegans is accompanied by a decline in the actin-cytoskeleton (Higuchi-Sanabria et al., 2018), we also found that in the shPab proteome and in aged TA muscles actin levels are reduced. Stabilization of actin filaments is regulated by actin binding proteins (Pollard, 2016, Vafiadaki, Arvanitis et al., 2014). Among those, CSRP3, PFN1, CFL1, ARP2/3 were found to be differentially expressed in shPab muscles (Olie et al., 2019). Stabilization of actin filaments in shPAB cell cultures, using actin overexpression, led to an increased fusion index in shPAB muscle bundles. Consistently, actin was shown to be expressed at the fusion site and was suggested to be essential for fusion (Stadler, Blattler et al., 2010). Nevertheless, actin overexpression did not completely restore myogenesis, as most shPAB MyHC expressing cells were smaller than in control fused cell cultures. This suggests that additional factors contribute to muscle atrophy in shPAB muscles and in aged muscles.

Expression of the actin-binding protein CSRP3 (also known as Muscle LIM protein (MLP)) was suggested to maintain the actin-cytoskeleton (Arber, Hunter et al., 1997, Hoffmann, Moreau et al., 2014). A role for CSRP3 in myogenesis was also demonstrated (Arvanitis, Vafiadaki et al., 2017, Vafiadaki et al., 2015). Here we found elevated CSRP3 expression in both shPab muscles and in aged TA muscles. Most prominent, CSRP3 expression highly correlated with MyHC-2A. MyHC-2A positive myofibers are more abundant in muscle atrophy and in shPab muscles (Riaz et al., 2016). This suggests that CSRP3 could mark atrophic myofibers. Increased CSRP3 levels were also found in different muscular dystrophies with muscle atrophy (Vafiadaki et al., 2015). The role of CSRP3 in atrophic myofibers is yet to be studied.

The expression of another structural protein, MURC/Cavin-4, was also consistent between aged TA muscles and shPab muscles. MURC is a member of the caveola gene family that regulates plasma membrane invagination and consequently plasma membrane-associated biological processes such as signaling, trafficking and mechanosensing (Bastiani et al., 2009). From those, the expression of both MURC and caveolin-3 (CAV3) is restricted to muscles (Kovtun, Tillu et al., 2015). MURC and Cav3 interact and form a complex at caveolae (Bastiani et al., 2009). Same as MURC, Cav3 protein was also found to have higher expression in the shPab proteome. Both proteins are involved in myogenesis (Bastiani et al., 2009, Tagawa et al., 2008). Loss-of-function CAV3 mutants may lead to different types of myopathies (Dewulf, Köster et al., 2019, Gonzalez Coraspe, Weis et al., 2018). Mutations in MURC were found in patients with dilated cardiomyopathy (Rodriguez, Ueyama et al., 2011). MURC function is not fully understood, but it is suggested to stabilize CAV3 and to regulate calcium trafficking across the membrane (Malette, Degrandmaison et al., 2019, Naito et al., 2015, Ogata, Naito et al., 2014). We show MURC expression in central nuclei and split myofibers, suggesting a role in regeneration. Consistently, increased MURC expression levels were also found in injury-induced muscle regeneration (Tagawa et al., 2008). MURC is essential for muscle cell fusion (Tagawa et al., 2008). Here we show that MURC localization at the plasma membrane in fused myotubes is regulated by the cytoskeleton. Destabilization of microtubules disrupts MURC localization at the plasma membrane, whereas stabilization of the cytoskeleton in shPAB myotubes, using actin overexpression, restores MURC’s plasma membrane localization. Moreover, MURC localization at the plasma membrane was also restored in the shPAB-actin-GFP muscle bundles. MURC expression was not found in the shPAB muscle bundles, but in the shPab muscles its expression at the myofibers’ boundaries was found as normal. This further suggests that MURC may play a role in aging-associated muscle regeneration.

To summarize, we showed a disordered cytoskeleton architecture in shPAB myoblasts and suggest a causative relation to reduced myogenesis. Furthermore, we found MURC as a novel marker for muscle regeneration. Stabilization of actin filaments, via actin overexpression, was beneficial for cell fusion in 2D or 3D models. The muscle bundle system seems beneficial for shPAB compared with the 2D cell culture system. The benefits of the muscle bundle model to study aging muscles should be investigated in future studies. The muscle bundle system seems to be beneficial to study PABPN1 regulation in muscle biology, as muscle bundles were formed from shPAB cells but the expression of sarcomeric and cytoskeletal genes was improper. This opens new opportunities to model muscle aging and muscle atrophy in vitro.

## Acknowledgments

This study was funded by the French Muscular Dystrophy Association (AFM-Téléthon) #26110 to VR.

The authors of this manuscript certify that they comply with the ethical guidelines for authorship and publishing in the Journal of Cachexia, Sarcopenia and Muscle. They have no conflict of interest.

## Supporting information

### Supplementary Tables

**Table S1.** Pearson correlation assessment of CSRP3 or MURC MFI with myosin heavy chain MFI. MFI of CSRP3 and MyHC-2A and -2B was measured from all myofibers in cross-sections of control and shPab muscles. A correlation between MFI’s was assessed with Pearson correlation coefficient. In total, over 1000 myofibers per genotype per an antibody-mix staining were included in the analysis.

|  | CSRP3 |  |
| --- | --- | --- |
| r (pvalue) | scram | shPab |
| MyhC-2A | 0.53<br>(9.8E-09) | 0.68 (0) |
| MyHC-2B | 0.14<br>(NS) | 0.003<br>(NS) |

### Supplementary Figures

**Fig S1.**
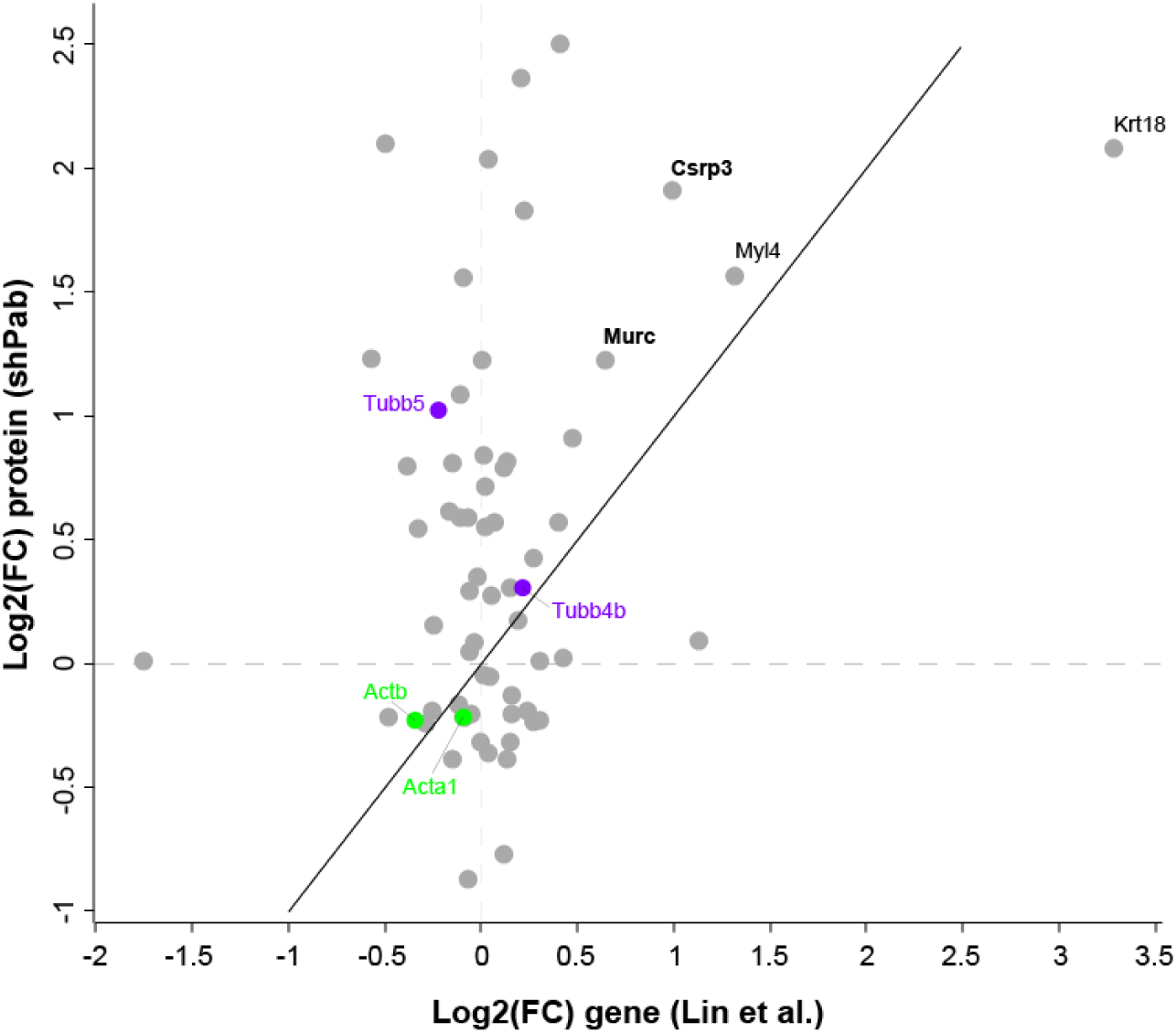
Fold-change comparison of muscle cytoskeletal genes in old mouse TA muscles to shPab mouse TA muscles. Scatter plot shows mRNA fold-changes (FC) in aged mice (Lin et al., 2018) to protein fold-changes in shPab muscles (Olie et al., 2019). Genes with FC>|1.5| in both studies are highlighted. The black line shows the diagonal. Actin genes/isoforms are denoted in green and tubulin in purple.

**Fig S2.**
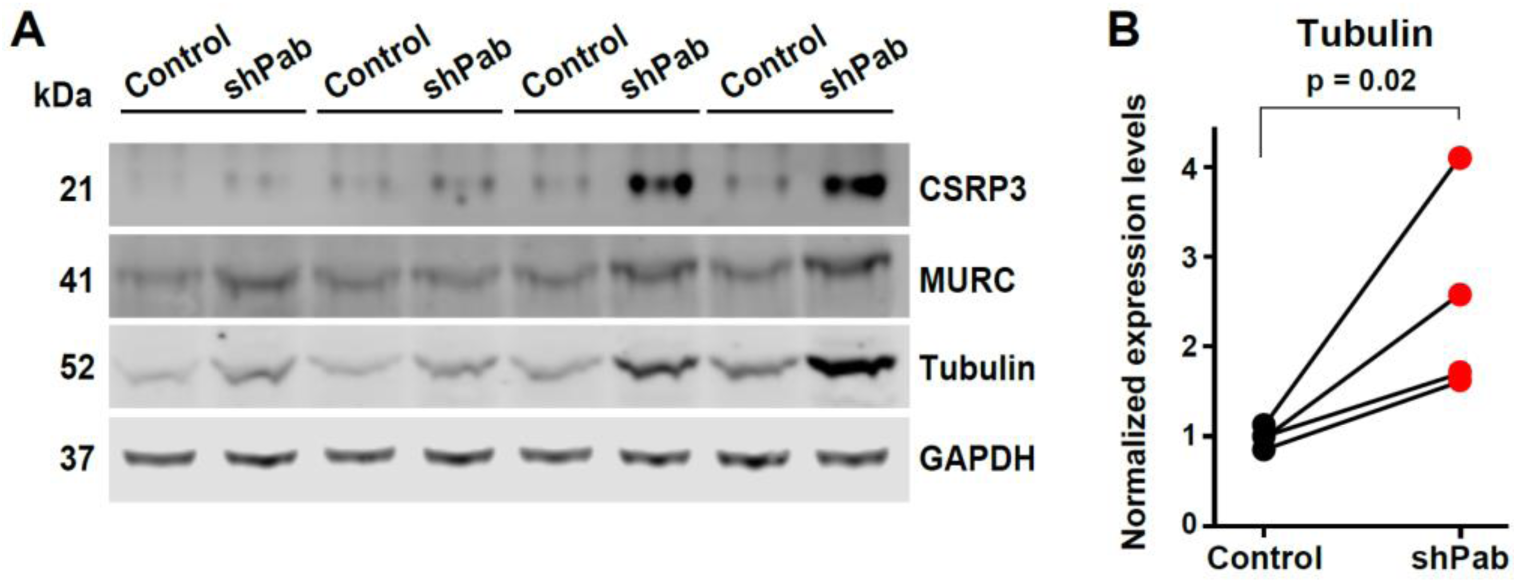
CSRP3, MURC and tubulin expression in paired tibialis anterior (control) versus shPab) muscles. **A.** Western blots show CSRP3, MURC and tubulin expression in control and shPab muscles. GAPDH was used as loading control. **B.** Western blot quantification. Paired dot-plot shows tubulin expression levels after normalization to loading control, N=4 mice. Statistical significance is assessed with a ratio-paired t-test.

**Fig S3.**
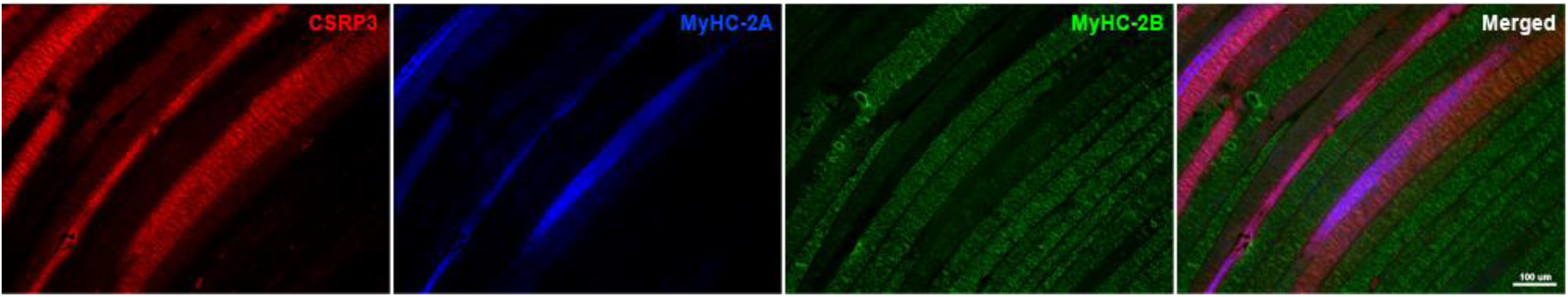
CSRP3 co-localization with MyHC-isoforms. Longitudinal sections of scram TA muscle showing MURC and MyHC-2A or 2B in TA muscle from control. Scale bar is 100 μm.

**Fig S4.**
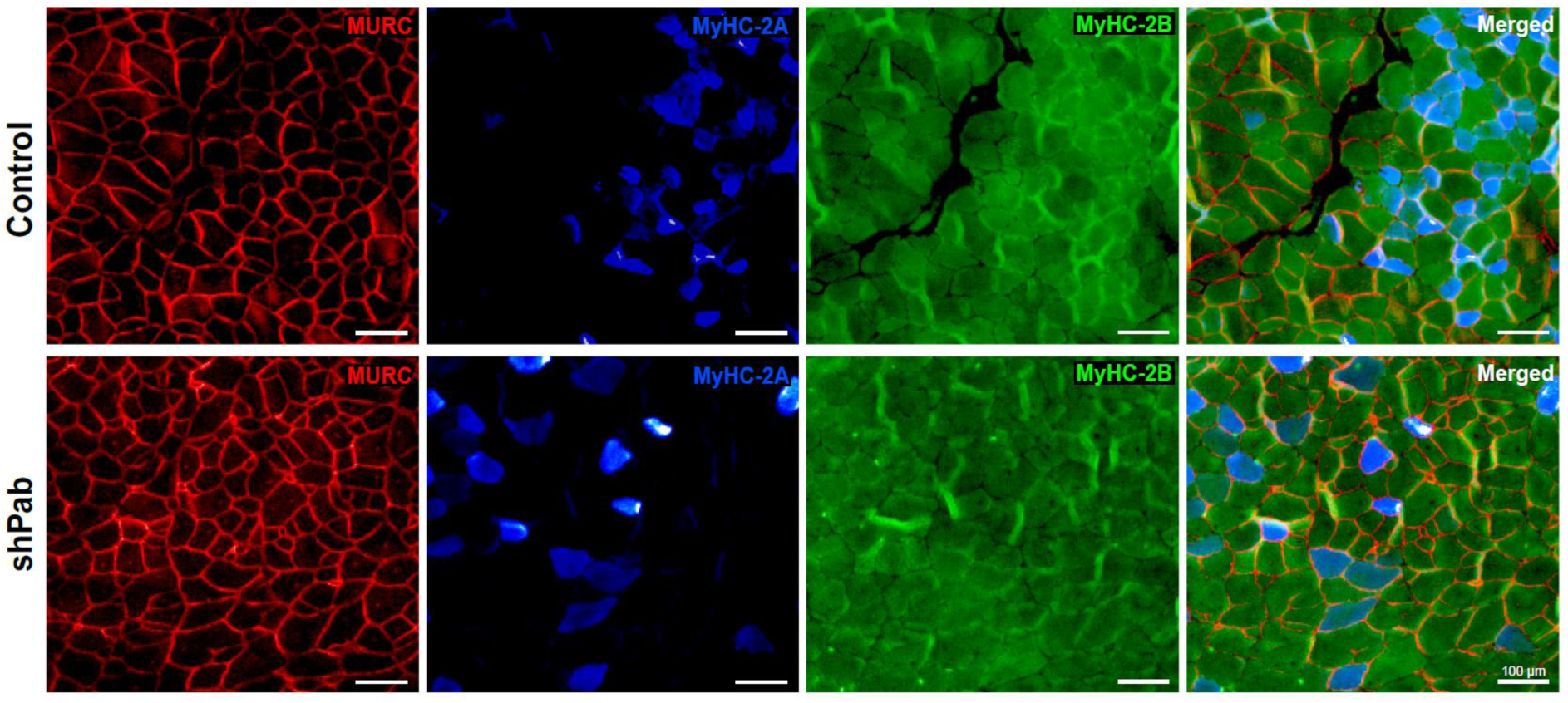
MURC co-localization with MyHC-isoforms. Representative images of MURC and MyHC-2A or 2B in control or shPab TA muscle cross-sections. Scale bar is 100 μm.

**Fig S5.**
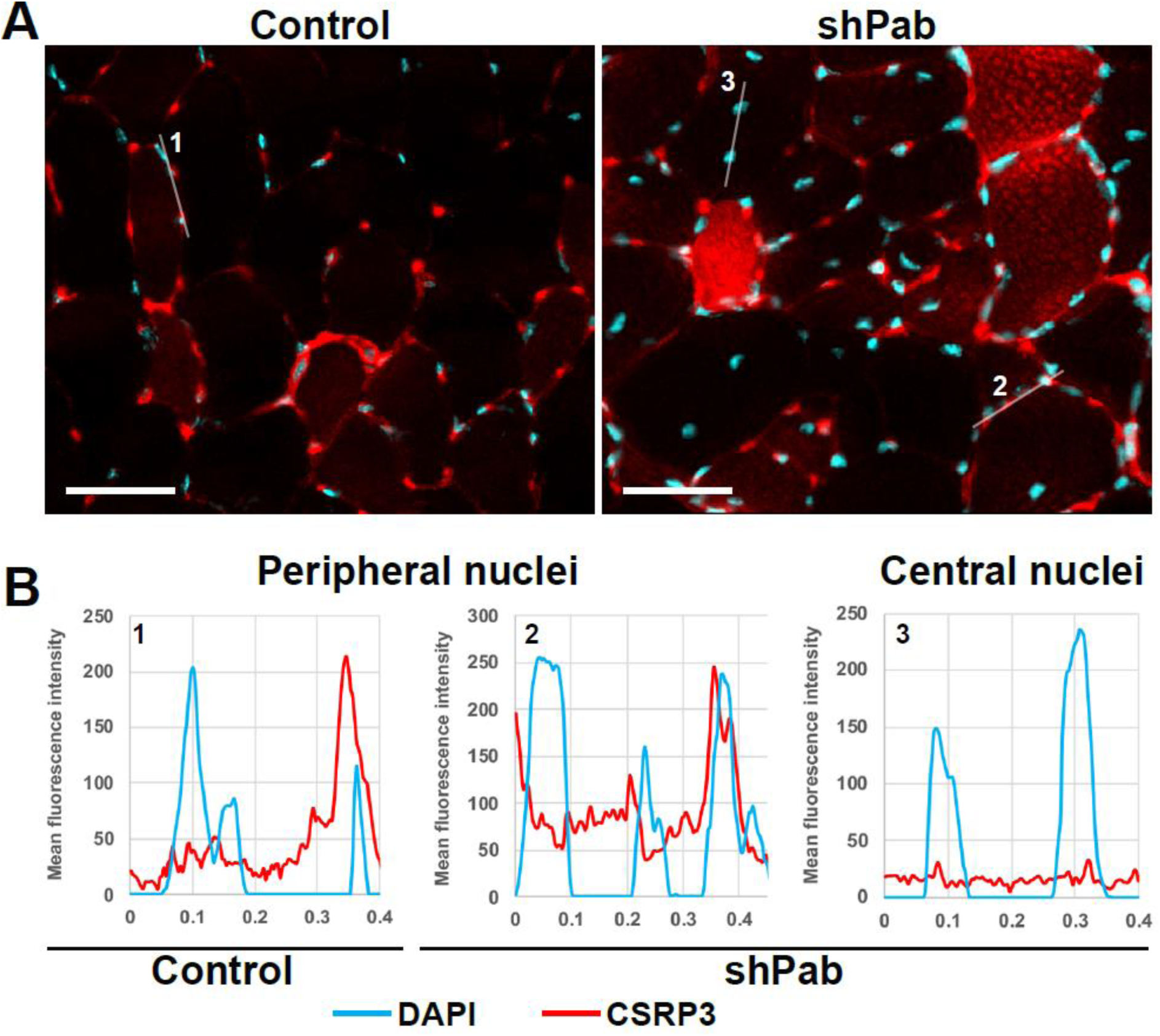
CSRP3 co-localization with myonuclei. **A.** Images of CSRP3 and DAPI in control or shPab TA muscle cross-sections. Scale bar is 50 μm. **B.** Intensity distribution plots show spatial overlap between CSRP3 and DAPI along the white lines, marked as 1 to 3. Two lines are from peripheral regions (#1 and #2), and #3 shows central nuclei. CSRP3 shows some overlap with peripheral myonuclei but not with central nuclei.

**Fig S6.**
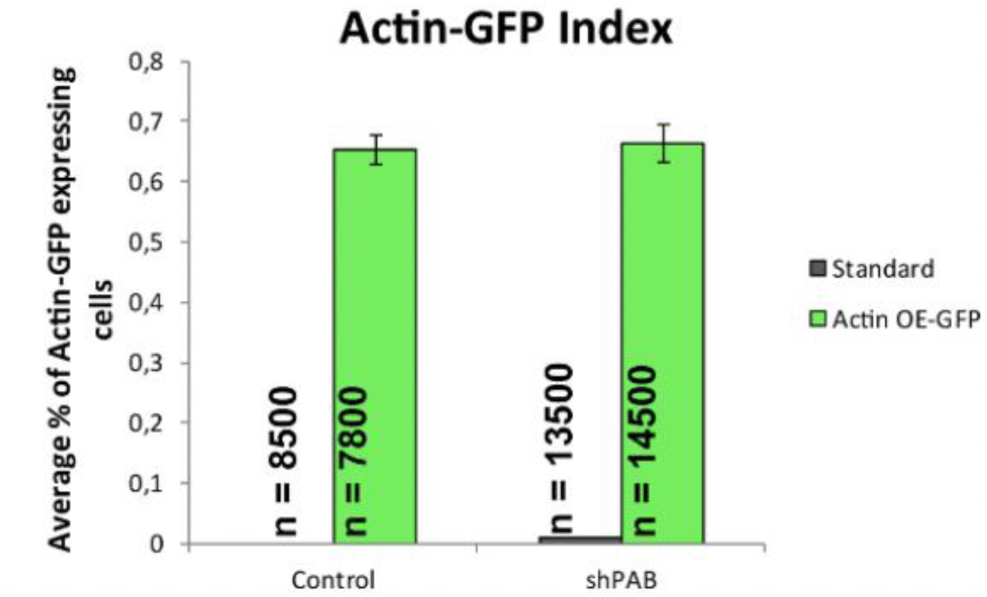
Percentage of GFP-positive cells after actin-GFP transduction of control or shPAB cell cultures. Parental cells were used as negative control for imaging in the GFP channel. N shows the average number of cells that were counted. Average and standard deviation are from three replicates.

